# Engineering liposomes with cell membrane proteins to disrupt melanosome transfer

**DOI:** 10.1101/2024.10.07.617008

**Authors:** Chunhuan Liu, Yuchun Liu, Changhu Xue, Cheng Yang, David A. Weitz, Kevin Jahnke

**Affiliations:** Key Laboratory of Synthetic and Biological Colloids, Ministry of Education, School of Chemical and Material Engineering, Jiangnan University, Wuxi 214122, China; School of Engineering and Applied Sciences, Harvard University, Cambridge, MA 02138; Key Laboratory of Marine Food Processing & Safety Control, School of Food Science and Engineering, Ocean University of China, Qingdao 266404, China; Department of Physics, Harvard University, Cambridge, MA 02138

## Abstract

Cells communicate by transporting vesicles and organelles, essential for maintaining cellular homeostasis. However, excessive vesicle transfer can cause several diseases and medical conditions like hyperpigmentation due to an unregulated intercellular transfer of melanosomes. Current treatments often focus on eliminating the compartment contents with drugs but can cause significant side effects. Here, we engineer liposomes with cell membrane proteins to directly disrupt intercellular transport without specialized therapeutics. We demonstrate this approach by reducing melanosome transfer from melanocytes to keratinocytes. To achieve this, we incorporate keratinocyte cell membrane proteins into liposomes using microfluidics, which can enhance uptake by melanocytes while reducing uptake by keratinocytes. We also show that these engineered liposomes reduce melanosome transfer because they attach to the surface of pigment globules, impeding pigment globule uptake by keratinocytes. Our findings provide an effective strategy for reducing melanosome transfer to treat hyperpigmentation and introduce a drug-free approach for regulating cellular communication via extracellular vesicles and organelles.

## Main

Cellular communication allows one cell to influence the behavior of another cell.^1–3^ To communicate, donor cells transport molecules, extracellular vesicles and organelles to recipient cells, activating signaling pathways that coordinate gene expression and thus drive cellular functions.^4–6^ Cellular communication via membrane-enclosed compartments such as exosomes, ectosomes and melanosomes is particularly important because they can carry genetic material from one cell to another.^7^ As a specific type of cellular communication, melanosome transfer drives the intercellular transport of natural pigments such as melanin in addition to genetic material. Thus, the transfer of melanosomes from melanocytes to keratinocytes governs mammalian pigmentation,^8, 9^ plays critical roles in the normal coloration formation of mammalian skin, hair and eyes, and regulates skin disease and cancer.^10–13^ An excess of melanin in keratinocytes leads to hyperpigmentation causing patches of skin to become darker than the surrounding skin.^14^ Typically, hyperpigmentation also includes melasma, a frequently acquired disorder, age spots and post-inflammatory hyperpigmentation types, which can affect the appearance, mental health, and social functioning of patients.^15, 16^ Many treatments have been focused on inhibiting the melanin production of cells by targeting tyrosinase or the cAMP pathway.^14^ However, the formation of melanin is an essential metabolic process of melanocytes, and an inhibition of this production will inevitably affect cell functionality and homeostasis. Therefore, melanin inhibition can cause serious side effects including chemical leukoderma.^17, 18^ In contrast to focusing on the melanin production itself, targeting the melanosome transfer process might be a particularly direct and promising strategy to prevent hyperpigmentation with fewer side effects. To achieve the transfer of melanin between cells, melanosomes are packaged into lipid vesicles called pigment globules, which are released into the extracellular space by melanocytes.^19, 20^ The pigment globules are then endocytosed by keratinocytes, after which they release the melanosomes, completing the melanosome transfer. To induce the endocytosis of melanosomes, pigment globules are recognized by the keratinocyte membrane through receptor-ligand interactions.^21^ If we knew the specific proteins and ligands involved in the endocytosis of pigment globules,^22^ we could specifically block these proteins and passivate the pigment globule surface,^23–26^ thus reducing the transfer of melanosomes. However, the receptors and ligands on the keratinocyte and melanocyte membrane that drive this interaction are unknown. Hence, there is a need to find alternate strategies to reduce melanosome transfer and to treat hyperpigmentation.

Here, we engineer biomimetic liposomes containing the whole cell membrane proteome of keratinocytes (HaCaT cell membrane proteins, HCMP), which can recognize and attach to pigment globules and melanocytes to reduce melanosome transfer. We fabricate biomimetic liposomes with HCMP (HCMP-liposomes) and show that the incorporation efficiency of HCMP is up to 90%. We also show that HCMP-liposomes have an enhanced uptake by human MNT-1 melanoma cells but a reduced uptake by HaCaT cells both in the single and co-culture model. We use this feature to significantly reduce the melanosome transport from MNT-1 to HaCaT cells by up to 70%, because HCMP-liposomes attach to the surface of pigment globules, impeding the uptake by HaCaT cells. Importantly, our approach of using the complete cell membrane protein extract to engineer liposomes is not limited to keratinocytes but could also be used more generally to regulate vesicle-based cellular communication based on extracellular vesicles to potentially prevent cancer metastasis^27, 28^ or neurodegenerative diseases.^29^

### HaCaT cell membrane proteins can be incorporated into liposomes

To engineer liposomes^30, 31^ that mimic the keratinocyte membrane,^32–34^ we extract HaCaT cell membrane proteins (HCMP) from immortalized human keratinocytes (HaCaT cells). We use the co-flow of a lipid-carrying and a protein-carrying solution on a microfluidic chip^35–37^ to form liposomes with HCMP (**Fig. 1a**). By injecting the aqueous HCMP solution into the outer channel and an ethanol solution with lecithin lipids and cholesterol into the inner channel, we produce uniform HCMP-liposomes with spherical shape and unilamellarity as shown in the cryogenic transmission electron microscopy (cryo-TEM) images (**Fig. 1b**). Scanning electron microscope (SEM) images reveal that both HCMP-liposomes and unmodified liposomes exhibit a spherical morphology (**Supplementary Fig. 1a, b**). We also find that we can control the liposome diameter by adjusting the flow rate ratio of the two phases and the total flow rate. By decreasing the ethanol-to-aqueous phase flow rate ratio (FRR) from 1:3 to 1:6, dynamic light scattering reveals that the liposome diameter decreases from 157 nm to 57 nm. An increase of the total flow rate (TFR) from 100 to 1000 *μ*L/min also decreases the liposome diameter by more than two-fold (**Supplementary Fig. 2a, b**). After the incorporation of HCMP into the liposome membrane, the mean diameter of liposomes increases by 20 nm indicating their successful incorporation (**Fig. 1c**). The polydispersity index (PDI) of HCMP-liposomes and unmodified liposomes is less than 0.2, indicating a high size homogeneity (**Fig. 1d**). We also perform zeta potential measurements to determine the effect of HCMP on the surface potential of liposomes. We find that HCMP-liposomes show a significant reduction of the surface potential to less than -20 mV, whereas unmodified liposomes have a zeta potential of -9 mV. Additionally, the zeta potential decreases with increasing HCMP-to-lipid ratios until it saturates at a HCMP to lipid ratio of 1:100 (**Fig. 1e**). The surface charge of HCMP-liposomes is similar to that of nanovesicles derived directly from HaCaT cell membranes with -24 ± 2 mV^38^, which indicates the successful incorporation and correct orientation of membrane proteins within HCMP-liposome bilayers. We also verify the successful incorporation of cell membrane proteins into the HCMP-liposomes with sodium dodecyl sulfate-polyacrylamide gel electrophoresis (SDS-PAGE) analysis (**Supplementary Fig. 3**).

**Figure 1.**
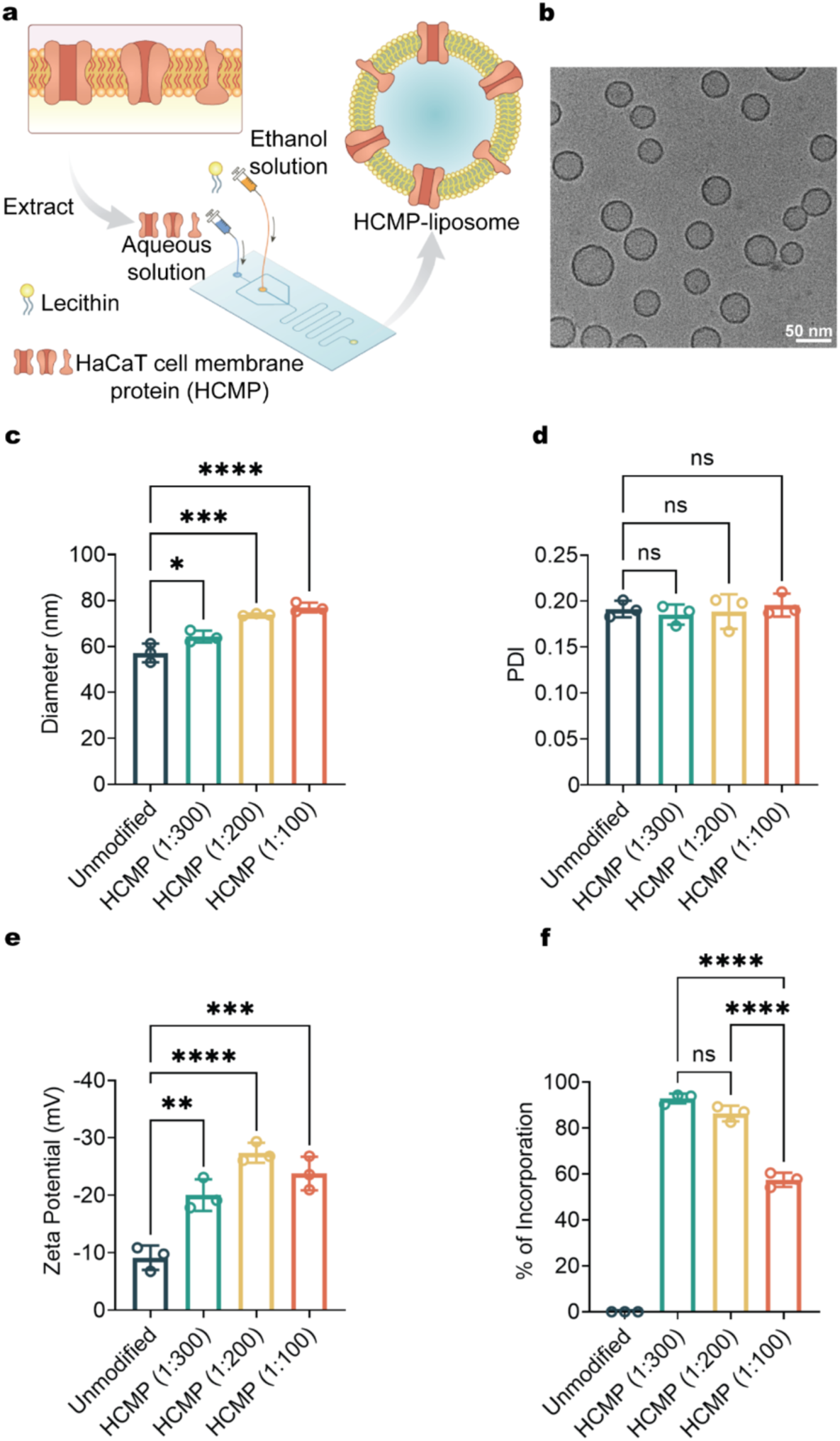
Fabrication and characterization of biomimetic liposomes with HaCaT cell membrane proteins (HCMP). **a** Schematic representation of HCMP-liposome formation with microfluidics. We inject the aqueous HCMP solution into the outer, and the ethanol solution containing lecithin and cholesterol into the inner channel. **b** Cryo-TEM images of HCMP-liposomes. Scale bar: 50 nm. **c**,**d** Hydrodynamic diameter (**c**) and polydispersity index (**d**) of unmodified and HCMP-liposomes at 1:300, 1:200, and 1:100 protein-to-lipid ratios. **e** Zeta potential of unmodified and HCMP-liposomes at 1:300, 1:200, and 1:100 protein-to-lipid ratios. **f** Evaluation of the HCMP incorporation efficiency into HCMP-liposomes at 1:300, 1:200, and 1:100 protein-to-lipid ratios as determined by a BCA Protein Assay Kit. Mean ± SD, n = 3.

To determine the incorporation efficiency of HCMP into liposomes, we purify them using dialysis and measure the protein concentration afterwards. We find that the incorporation efficiency is as high as 93% for HCMP-liposomes (1:300) (**Fig. 1f**). From this, we approximate that about 15% of the HCMP-liposome surface area is covered by membrane proteins (**Supplementary Note)**. Although HCMP-liposomes have the highest HCMP density at an HCMP-to-lipid ratio of 1:100, the HCMP incorporation efficiency is the lowest, indicating the saturation of the membrane with HCMP and the presence of HCMP in the surrounding solution. Hence, we set the HCMP-to-lipid ratio to 1:200 for the following experiments.

### HCMP-liposomes have an enhanced uptake by MNT-1 cells

To investigate whether HCMP-liposomes can specifically interact with target cells, we select the darkly pigmented melanoma cell line MNT-1.^39^ MNT-1 cell membrane ligands are known to be recognized by HCMP.^9, 40^ We also select HaCaT cells as a control because they do not have ligands which are recognized by HCMP.

To investigate the uptake of HCMP-liposomes by cells, we incubate HaCaT and MNT-1 cells separately with 500 *μ*g/mL rhodamine B-labeled liposomes. This lipid concentration ensures a high cell viability (**Supplementary Fig. 4a-c**). From the confocal images, we observe that cells incubated with HCMP-liposomes show a reduced lipid fluorescence intensity compared to unmodified liposomes. By contrast, MNT-1 cells incubated with HCMP-liposomes show an increased lipid fluorescence compared to unmodified liposomes (**Fig. 2a**). We quantify the average lipid fluorescence signal in the confocal images and find that the average fluorescence intensity in HaCaT cells incubated with HCMP-liposomes is less than half of that of unmodified liposomes (**Fig. 2b**). However, in MNT-1 cells, the average fluorescence intensity in cells incubated with HCMP-liposomes is 1.5-fold higher than that of unmodified liposomes (**Fig. 2c and d**). We also analyze the mean fluorescence intensity (MFI) per cell with flow cytometry, which provides additional evidence that HCMP-liposomes exhibit a significantly reduced uptake in HaCaT cells (**Fig. 2e**) and an increased uptake by MNT-1 cells (**Fig. 2f**). To confirm whether the HCMP-liposomes can actively interact with other melanocytes, we choose A375 as a second cell line isolated from human malignant melanoma and test its uptake of liposomes. Similar to MNT-1 cells, both the confocal and flow cytometry data demonstrate a significantly higher cell uptake by A375 cells for HCMP-liposomes compared to unmodified liposomes (**Supplementary Fig. 5** and **Fig. 6**). Overall, these results show that HCMP-liposomes can increase the uptake by melanocytes while reducing the uptake by HaCaT cells.

**Figure 2.**
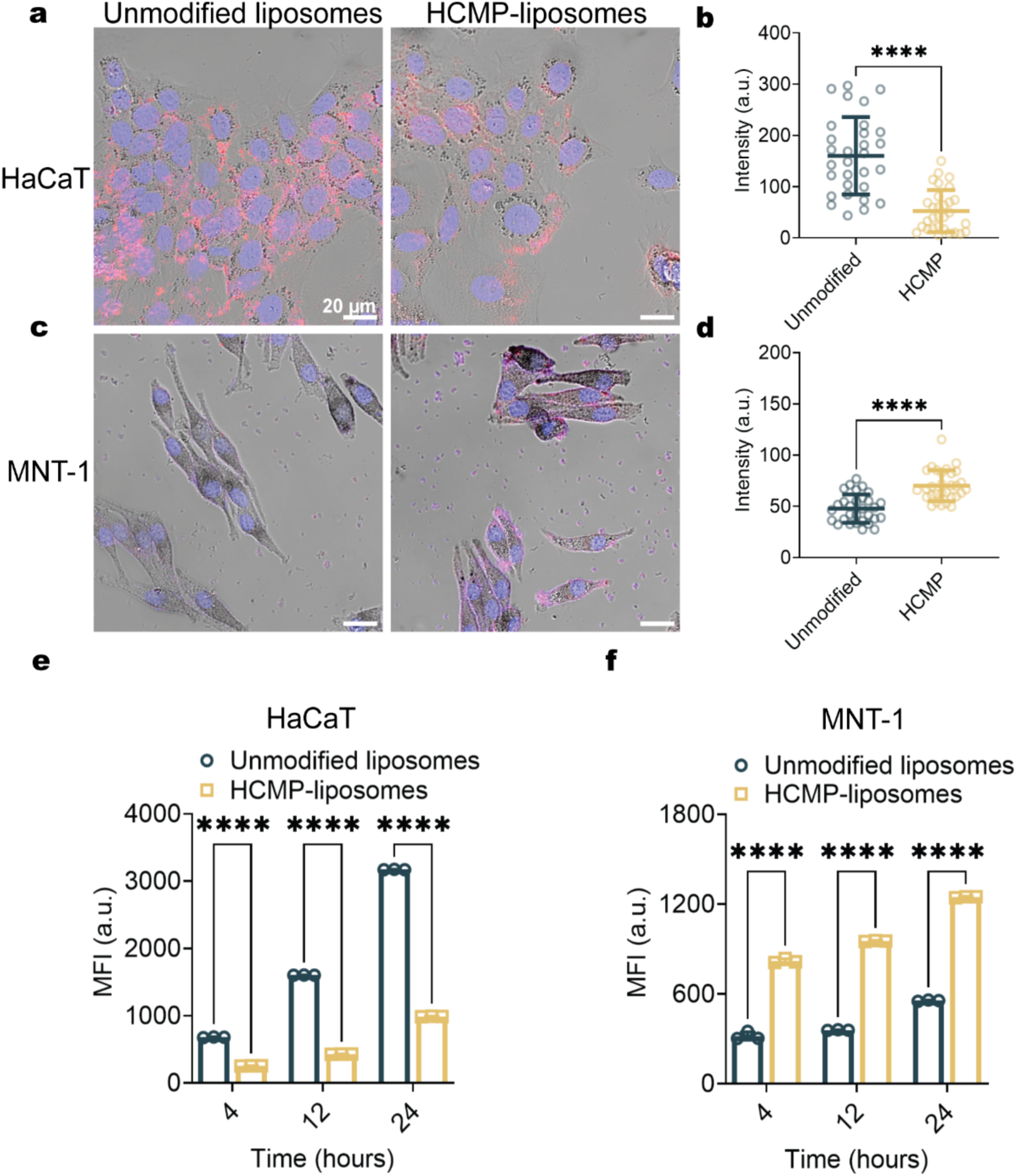
HCMP-liposomes show enhanced uptake by MNT-1 cells and a reduced uptake by HaCaT cells. **a** Confocal images showing the HaCaT (blue, DAPI-stained; *λex* = 405 nm) cell uptake of unmodified and HCMP-liposomes (red, rhodamine B-labeled; *λex* = 560 nm) after 24 h. **b** Fluorescence intensity analysis of liposomes uptake by HaCaT cells, Mean ± SD, n = 30. Scale bar: 20 *µ*m. **c** Confocal images showing the MNT-1 (blue, DAPI-stained; *λex* = 405 nm) cell uptake of unmodified and HCMP-liposomes (red, rhodamine B-labeled; *λex* = 560 nm) after 24 h. **d** Fluorescence intensity analysis of liposomes uptake by MNT-1 cells, Mean ± SD, n = 30. Scale bar: 20 *µ*m. **e**,**f** Mean fluorescence intensity (MFI) obtained from flow cytometry showing the uptake of unmodified and HCMP-liposomes by HaCaT (**e**) and MNT-1 (**f**) cells after 4, 12 and 24 h. Mean ± SD, n = 3 for 30,000 events, respectively.

### HCMP-liposomes are selectively taken up by MNT-1 cells

To mimic the epidermal microenvironment and verify the selective uptake of HCMP-liposomes by melanocytes, we establish a co-culture model of HaCaT and MNT-1 cells (**Fig. 3a**). To distinguish between cell types, we label HaCaT cells fluorescently with Cell-Tracker Green. Confocal imaging reveals that the HaCaT cells exhibit a bright and uniform green fluorescence throughout the cytoplasm, whereas MNT-1 cells do not show any green fluorescence (**Fig. 3b**). After the addition of liposomes, we observe that HaCaT cells exhibit a lower lipid fluorescence signal for HCMP-liposomes compared to unmodified liposomes. In contrast, MNT-1 cells show higher lipid fluorescence signals of HCMP-liposomes compared to unmodified liposomes (**Fig. 3b and c**), which is consistent with the results from the single culture model.

**Figure 3.**
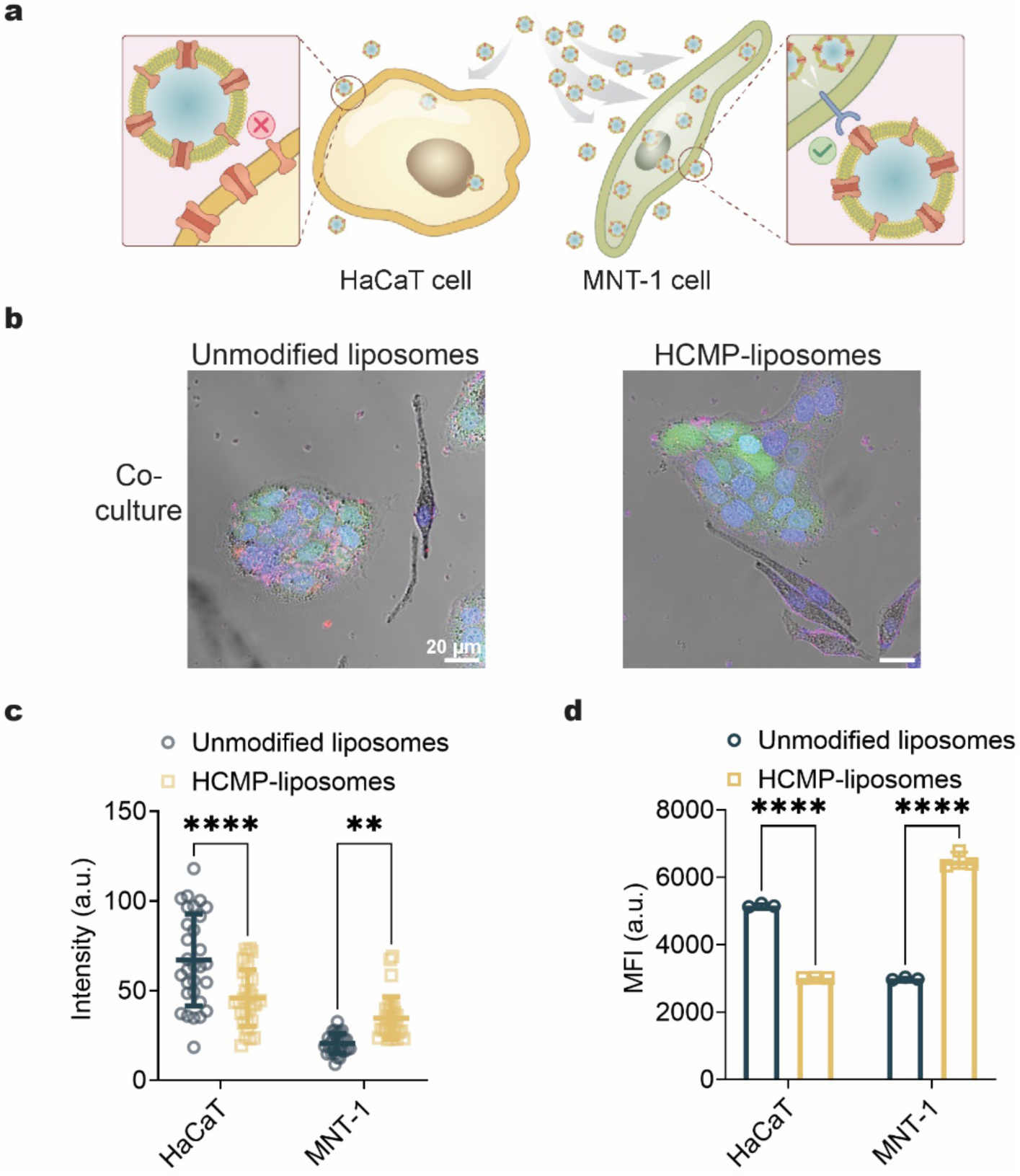
A co-culture model reveals that HCMP-liposomes have an enhanced uptake by MNT-1 cells compared to HaCaT cells. **a** Schematic representation of the selective uptake of HCMP-liposomes by MNT-1 cells. **b** Confocal images showing the uptake of unmodified and HCMP-liposomes (red, rhodamine B-labeled, *λex* = 560 nm) by HaCaT (green, Cell-Tracker Green CMFDA-labeled, *λex* = 488 nm, blue: DAPI-stained, *λex* = 405 nm) and MNT-1 cells (blue: DAPI-stained, *λex* = 405 nm) after 24 h. **c** Fluorescence intensity analysis of liposomes uptake by HaCaT and MNT-1 cells. Mean ± SD, n = 30. Scale bar: 20 *µ*m. **d** Mean fluorescence intensity (MFI) obtained from flow cytometry showing the uptake of unmodified and HCMP-liposomes by HaCaT and MNT-1 cells after 24 h. Mean ± SD, n = 3 for 30,000 events, respectively.

We also use flow cytometry to quantify the preferential cell uptake in the co-culture model. To distinguish the two cell types, we also label MNT-1 cells with Cell-Tracker Deep Red (**Supplementary Fig. 7**). HaCaT cells treated with HCMP-liposomes exhibit a 40 % reduction in the mean fluorescence intensity (MFI) compared to HaCaT cells treated with unmodified liposomes. By contrast, MNT-1 cells treated with HCMP-liposomes show a 120 % increase in the MFI compared to unmodified liposomes. This enhanced uptake is driven by specific binding interactions with receptors on the MNT-1 membrane and HCMP, promoting the internalization of HCMP-liposomes by MNT-1 cells.

### HCMP-liposomes reduce melanosome transfer to HaCaT cells

MNT-1 cells continuously produce and release melanosomes^8^, which are then phagocytosed by HaCaT cells (**Fig. 4a**). In this process, melanosomes display specific binding and uptake by HaCaT cells due to receptor-mediated interactions. We test if we can use the interactions of HCMP-liposomes with MNT-1 cell membranes to reduce the melanosome uptake by HaCaT cells.

**Figure 4.**
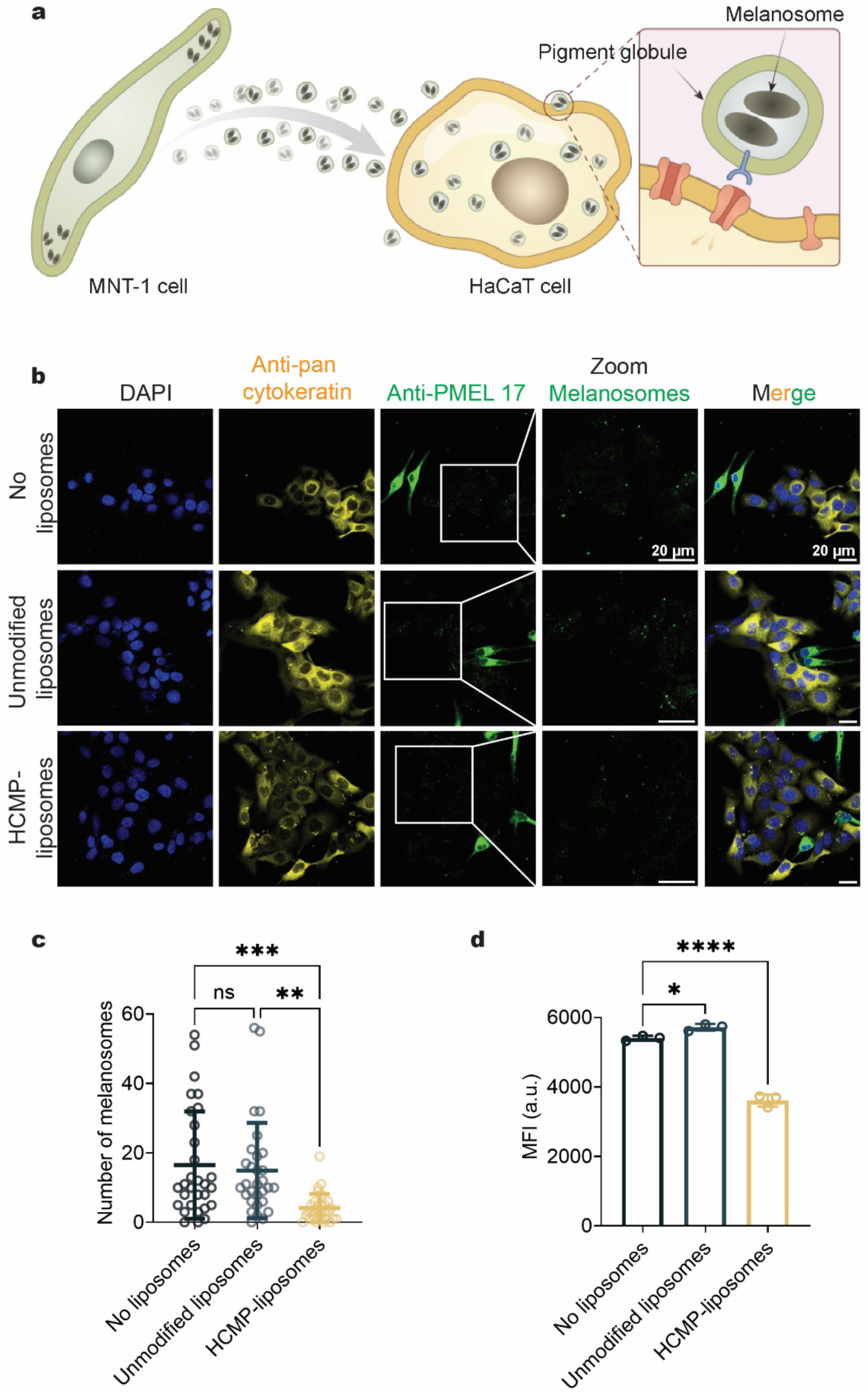
HCMP-liposomes reduce melanosome transfer between HaCaT and MNT-1 cells. **a** Schematic representation of melanosome transfer process. **b** Confocal images of co-cultured HaCaT and MNT-1 cells incubated without liposomes, with unmodified liposomes and with HCMP-liposomes after 24 h. Scale bar: 20 *µ*m. Blue: nuclei stained with DAPI (*λex* = 405 nm); Yellow: HaCaT cells immunostaining with Anti-pan cytokeratin (*λex* = 640 nm); Green: MNT-1 cells and melanosomes immunostaining with Anti-PMEL 17 (*λex* = 488 nm). **c** Transferred melanosomes per HaCaT cell. Values are expressed as the number of PMEL-17-positive spots per HaCaT cell for each group. Mean ± SD, n = 30. **d** Mean fluorescence intensity (MFI) measured with flow cytometry showing melanosomes transferred to HaCaT cells after 24 h in absence of liposomes, after the addition of unmodified or HCMP-liposomes. Mean ± SD, n = 3 for 30,000 events, respectively.

To ensure a high rate of melanosome transport, we use a co-culture model with a HaCaT to MNT-1 cell ratio of 1:5 (**Supplementary Fig. 8a and 8b**). In the co-culture model, we evaluate the melanosome transfer by tracking the number of fluorescently-labeled melanosomes inside HaCaT cells with confocal microscopy.^41^ We observe a reduction of melanosomes within HaCaT cells in the group treated with HCMP-liposomes compared to either adding no liposomes or unmodified liposomes (**Fig. 4b**). To quantify the melanosome transfer efficiency into HaCaT cells, we count the number of melanosome spots in their cytoplasm (**Fig. 4c**). We find that the number of melanosome spots decreases 4-fold in groups treated with HCMP-liposomes but does not decrease in cells treated with unmodified liposomes.

We also confirm these results with flow cytometry^42, 43^ (**Supplementary Fig. 9**), where the MFI of melanosomes in HaCaT cells treated with HCMP-liposomes shows a significant reduction compared to groups without liposomes or unmodified liposomes (**Fig. 4d**). Our results validate the hypothesis that HCMP-liposomes exert a significant inhibitory effect on melanosome transfer, whereas unmodified liposomes do not.

### HCMP-liposomes passivate the pigment globule surface

To reveal the mechanism, by which HCMP-liposomes reduce melanosome transfer, we label the melanosomes of MNT-1 cells with the antibody PMEL 17 shown in cyan. We find that the pigment globules with melanosomes are secreted into the extracellular space (**Supplementary Fig. 10, Supplementary Fig. 11**). This is consistent with the model of the phagocytosis of melanosome-rich pigment globules.^44^ The model postulates that pigment globules containing melanosomes are released into the extracellular space from melanocytes and then ingested by keratinocytes^19, 21^. We speculate that the pigment globules interact with HCMP-liposomes when released into the extracellular space, leading to a reduction of melanosome transfer (**Fig. 5a**).

**Figure 5.**
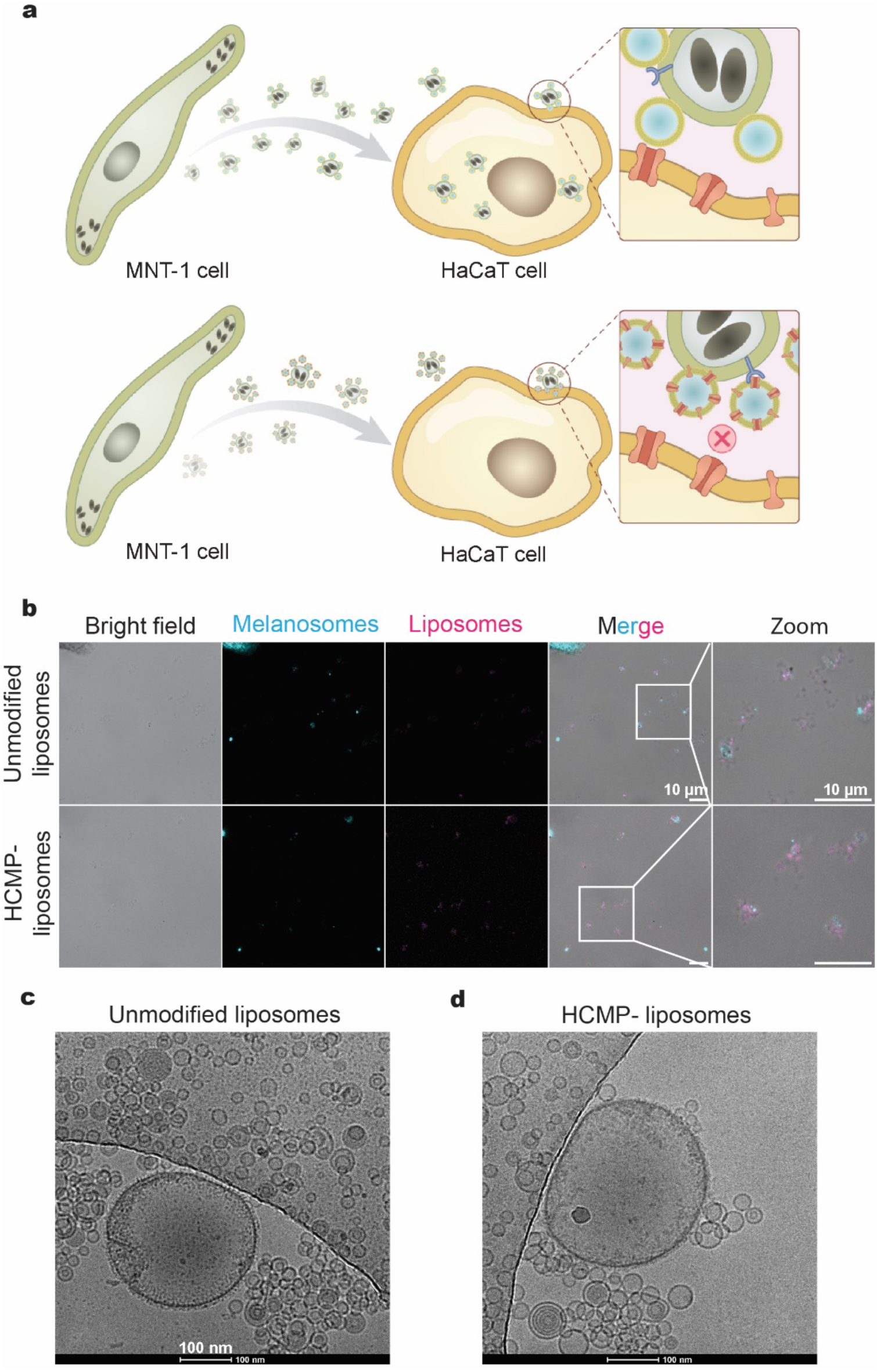
Proposed mechanism of the disruption of melanosome transfer by with HCMP-liposomes. **a** Schematic representation of the mechanism by which HCMP-liposomes reduce melanosome transfer. **b** Confocal images of melanosomes (cyan, immunostained with Anti-PMEL 17; *λex* = 488 nm) with unmodified liposomes and HCMP-liposomes (purple, rhodamine B-labeled; *λex* = 560 nm) after 24 h. Scale bar: 10 *µ*m. **c,d** Cryo-TEM images of extracted melanosomes with unmodified liposomes (**c**) and HCMP-liposomes (**d**). Scale bar: 100 nm.

To investigate the interaction between HCMP-liposomes and pigment globules in the extracellular space, we label liposomes with rhodamine B shown in purple, and find that pigment globules outside MNT-1 cells are almost entirely surrounded by liposomes (**Fig. 5b**). To investigate the interaction between pigment globules and HCMP-liposomes in the absence of cells, we isolate the pigment globules from the culture medium of MNT-1 cells, and separately incubate them with rhodamine B-labeled liposomes. We find a striking co-localization between unmodified and HCMP-liposomes and isolated pigment globules (**Supplementary Fig. 12**). The unmodified liposomes form large fluorescence spots indicating the aggregation of liposomes. We do not find any aggregation of HCMP-liposomes in the cell medium in the absence of pigment globules (**Supplementary Fig. 13**). We additionally confirm the interaction of HCMP-liposomes with pigment globules with cryo-transmission electron microscopy (TEM). The isolated pigment globule is about 200 nm and shows a clear membrane structure (**Supplementary Fig. 14**), which is surrounded by liposomes (**Fig. 5c, d**). This indicates that HCMP-liposomes anchor to the pigment globules. By contrast, unmodified liposomes tend to aggregate. Thus, these results indicate that HCMP-liposomes can anchor onto the surface of pigment globules and thereby alter their recognition by HaCaT cells during cellular phagocytosis of pigment globules. Instead of displaying MNT-1 cell membrane proteins and receptors, pigment globules are covered by HaCaT cell membrane proteins, which reduces the uptake of pigment globules and melanosomes.

## Conclusion

We engineer liposomes with HCMP and show that HCMP-liposomes specifically interact with MNT-1 cells but not with HaCaT cells. We use this feature to disrupt the melanosome transport from MNT-1 to HaCaT cells. Our findings establish an effective strategy for reducing melanosome transfer and to treat hyperpigmentation without any synthesized therapeutics. They also highlight that HCMP-liposomes can offer a promising avenue for targeted therapeutic interventions in pigmentary disorders and related skin conditions. This strategy is expected to be useful for clinical application after further assessments in *in vivo* studies.

Additionally, we find that HCMP-liposomes have a specific recognition effect on melanocytes leading to a significant increase of their uptake by melanocytes. Thus, the incorporation of HCMP could be an important strategy for the targeted delivery of therapeutics through liposomes to melanoma, the most dangerous type of skin cancer. Our findings also introduce a general materials-based approach for regulating cellular communication pathways between cell types, when the targeted membrane proteins are unknown. Liposomes engineered with proteins from other cell types could be used to specifically target cell membrane-based vesicles, including exosomes and ectosomes, and modulate their recognition by recipient cells. Thus, our method could also form the basis of new treatment options to prevent cancer metastasis or neurodegenerative diseases and enables new perspectives for medical research and clinical practice.

## Materials and methods

### Materials

The microfluidic poly (dimethylsiloxane) (PDMS) chip (the channel depth is 100 *μ*m and the width is 200 *μ*m, **Fig. 1a**) was purchased from Jianmi Zhikong Technology Co, Ltd., Wuhan, China. 1-Palmitoyl-2-linoleoyl-sn-glycero-3-phosphocholine (Soybean phospholipid, purity > 98%) and Rhodamine B 1, 2-dihexadecanoyl-sn-glycero-3-phosphoethanolamine (triethylammonium salt) (Rhodamine DHPE) were purchased from Aladdin Biochemical Technology Co., Ltd., (Shanghai, China). Cholesterol and HPLC-grade solvents were purchased from Sinopharm Chemical Reagent Co., Ltd., (Shanghai, China). The HaCaT cell line was purchased from the BeNa Culture Collection (Beijing China). The MNT-1 cell line was purchased from the Zhejiang Meisen Cell Technology Co., Ltd. (Hangzhou, China). 1,2-dioleoyl-sn-glycero-3-phosphocholine (DOPC) and 1,2-dioleoyl-sn-glycero-3-phosphoethanolamine-N-(lissamine rhodamine B sulfonyl) (ammonium salt) (18:1 Liss Rhod PE) in chloroform form were purchased from Avanti polar lipids (USA). The A375 cell line was purchased from ATCC (USA).

### Cell Membrane Protein Extraction and Quantification

HaCaT cell membrane proteins (HCMP) were extracted from HaCaT cells using a Membrane and Cytosol Protein Extraction Kit (P0033, Beyotime, Shanghai, China) according to the manufacturer’s protocol. Quantification of extracted membrane proteins was performed using a BCA Protein Assay Kit (P0012, Beyotime, Shanghai, China) according to the manufacturer’s protocol. Absorbance was measured at 562 nm at a multi-mode microplate reader (Molecular Devices, Sunnyvale, USA), and the membrane protein concentration was determined using a standard curve. Extracted HaCaT cell membrane protein supernatants were stored with phenylmethanesulfonyl fluoride (Beyotime, Shanghai, China) at -80 ℃ until use.

### Identification of cell membrane proteins

The composition of total protein, membrane protein and cytosol protein of HaCaT cells and protein composition of HCMP-liposomes with different membrane protein concentrations were characterized by the sodium dodecyl sulfate-polyacrylamide gel electrophoresis (SDS-PAGE) method.

### Unmodified liposomes and HCMP-liposomes preparation and purification

Liposomes were synthesized using a microfluidic chip (**Fig. 1a**). Soybean phospholipid and cholesterol (7:3 molar ratio) were dissolved in ethanol at a final lipid concentration of 10 mg/mL and loaded in the inlet of the organic phase (**Fig. 1a**, ethanol solution syringe). The aqueous buffer and the ethanol solution of lipids were mixed between the inlet streams at different total flow rates (TFR) and flow rate ratios (FRR) to characterize liposomes of size, polydispersity index (PDI) and zeta potential. After that, unmodified liposomes were dialyzed overnight using 300 kDa dialysis bags (Spectrum Labs, Los Angeles, USA) at 4 ℃ in 1 × PBS (1:1000 v/v, Cytiva, Logan, USA), with one buffer change every 1 h. After dialysis, unmodified liposomes were collected and filtered using 0.22 *μ*m PES syringe filters (Titan, Shanghai, China). HCMP-liposomes were prepared using the same method but with HCMP redissolved in aqueous buffer (1 × PBS) at a protein-to-lipid concentration ratio of 1:100, 1:200, or 1:300 (w/w) and loaded in the aqueous phase inlet. HCMP-liposomes were assembled with the microfluidic chip using the following parameters: 6:1 FRR, 1 mL min^-1^ TFR, initial waste = 0.1 mL, final waste = 0.05 mL (**Supplementary Fig. 1**). HCMP-liposomes were then dialyzed overnight using 300 kDa dialysis bags (Spectrum Labs, Los Angeles, USA) at 4 °C in 1 × PBS (1:1000 v/v), with one buffer change after 1 h. After dialysis, liposomes were collected and filtered using 0.22 *μ*m PES syringe filters (Titan, Shanghai, China).

### Characterization of Liposome Size, Polydispersity Index (PDI), Zeta Potential, and membrane proteins’ incorporation

Dynamic light scattering (DLS) (Brookhaven, Holtsville, USA) was used to determine the size, PDI and zeta potential of liposomes. 300 *μ*L sample was diluted with 2.7 mL of 1 × PBS. The diluted samples were added in polystyrene cuvettes (Sheepurk, Wuxi, China) for size and PDI measurements. For each sample, a total of three runs with 20 measurements/run were performed and the average of these three runs was reported. For the zeta potential measurements, 20 *μ*L of the sample was diluted with 1.98 mL 1 × PBS. Diluted samples were transferred into polystyrene cuvettes. For each measurement, three runs with 30 measurements/run were performed; the average of these three runs was reported.

The amount of protein incorporated into HCMP-liposomes was determined from the difference between the amount of protein initially added to the formulation and the amount that was not incorporated into HCMP-liposomes. Briefly, after mixing in the microfluidic chips, the samples were dialyzed by using 3000 Da dialysis bags (Spectrum Labs, Los Angeles, USA) to remove lipids in order to avoid the interference of lipids with the BCA assay. After dialysis, the samples were ultracentrifuged at 8000 × g for 20 min using a 300 kDa Ultrafiltration centrifuge tube (Pall, New York, USA) to separate the liposomes from the unincorporated membrane proteins. The supernatant containing the membrane proteins was analyzed through a BCA Protein Assay Kit (Beyotime, Shanghai, China) to quantify the amount of membrane proteins not incorporated into liposomes, and the amount of membrane proteins incorporated into liposomes was calculated using the following equation.

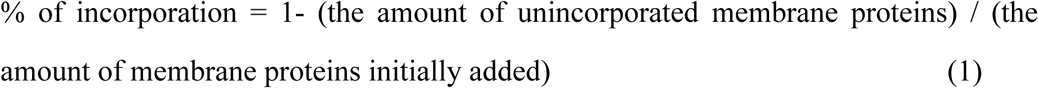

### Scanning electron microscopy (SEM) and cryogenic transmission electron microscopy (Cryo-TEM) of liposomes

For SEM, freshly prepared liposomes were rapidly frozen in liquid nitrogen with 50 mM trehalose, and then the samples were lyophilized in a vacuum overnight using freeze dryer (SCIENTZ, Ningbo, China). After that, the lyophilized dry powder was applied to a carbon double-sided tape (NISSHIN-EM Co., Ltd., Tokyo, Japan), covered by a thin layer of gold, and visualized in a S-4800 SEM (Hitachi, Tokyo, Japan).

For Cryo-TEM, the experiments were performed with thin films of sample solution of unmodified liposomes and HCMP-liposomes (5 *µ*L) transferred to a lacey supported grid in room temperature (Lacey Formvar/Carbon, 200 mesh, Cu; Ted Pella, Inc.). The thin solution films were prepared under controlled temperature and humidity conditions (97-99%) within a custom–built environmental chamber. The excess liquid was blotted with filter paper for 2-3 seconds, and the thin solution films were rapidly vitrified by plunging them into liquid ethane (cooled by liquid nitrogen) at its freezing point. The grid was transferred, on a Gatan 626 cryo-holder, using a cryo-transfer device and transferred to the Talos F200C TEM instrument (200 kV). Direct imaging was carried out at a temperature of approximately -175 °C and with a 200 kV accelerating voltage, using the images acquired with a SC 1000 CCD camera (Gatan, Inc., USA). The data were analyzed using Digital Micrograph software (Gatan, Inc., USA) and ImageJ (National Institutes of Health).

### Cell Culture

HaCaT cells and A375 cells were cultured in DMEM (Gibco, Grand Island, USA) supplemented with 10% Fetal Bovine Serum (FBS, Newserum, Christchurch, New Zealand) and 1% Penicillin/Streptomycin (Gibco, Grand Island, USA). MNT-1 cells were cultured in DMEM supplemented with 20% FBS, 1% NEAA (Gibco) and 1% Penicillin/Streptomycin. For co-culture models, HaCaT and MNT-1 cells with the ratio of 1:1, 1:2, 1:5 and 1:10, were seeded in a 6-well plate (Titan, Shanghai, China) at a cell density of 5 × 10^5^ cells/well. The co-culture media was composed of HaCaT medium and MNT-1 medium at the ratio of 1:1, 2:1, 5:1, and 10:1, respectively. All cells were cultured in a humidified incubator (ThermoFisher Scientific, Waltham, USA) with 95% air and 5% CO2 at 37 ℃.

### Cell Cytotoxicity

HaCaT, MNT-1 cells and co-culture models were seeded in a 96-well plate at a density of 10^4^ cells per well in respective medium, and cultured at 37 °C under 5% CO2 for 24 h respectively. A cell-free group was used as blank. After 24 h, the medium was discarded, and the cells were washed twice with PBS. For the experimental groups, the cells were incubated with a medium containing the tested liposomes, and the concentration of liposomes was set to 0.1, 0.5, 1 mg/mL; for the control and blank group, only medium was added. After incubation for 24 h, 10 *μ*L of CCK-8 solution (Beyotime, Shanghai, China) was added to each well and then incubated in the cell incubator for 1h. After incubation, the absorbance of the test group was monitored at 450 nm using a microplate reader. The optical density (OD) values of the experimental group, blank group, and control group were recorded as ODe, ODb, and ODc, respectively, and the cell viability was calculated using the following equation.

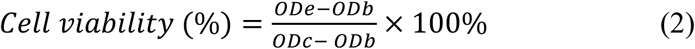

### Cell uptake in HaCaT and MNT-1 cells

HaCaT and MNT-1 cells were used to evaluate the cell uptake of unmodified liposomes and HCMP-liposomes. Liposomes were labeled with rhodamine B by adding 0.2 mol% Rhodamine DHPE in the organic phase, followed by preparation procedures as described above. **Confocal observation experiments**: HaCaT and MNT-1 cells were cultured in a culture dish (Wuxi NEST Biotech Co., Ltd., Wuxi, China) at a density of 1 × 10^5^ per well for observation with a Zeiss LSM880 with Airyscan confocal laser scanning microscope (CLSM, ZEISS, Oberkochen, Germany), the 20X objective and 100X oil-immersion objective were used. After 24 h, the cells were washed twice with 1 × PBS, and incubated with rhodamine B-labeled liposomes diluted with respective medium to a liposome concentration of 0.5 mg/mL for 24 h at 37 ℃. After treatment, the cells were fixed with 4% Paraformaldehyde Fix Solution (Beyotime, Shanghai, China) for 10 min, stained with DAPI Staining Solution (Beyotime, Shanghai, China) for 4 min, and then imaged via CLSM (Ex/Em: DAPI, 364/454 nm; rhodamine B, 560/580 nm). ImageJ (National Institutes of Health) was used to quantify the relative fluorescence intensity of rhodamine B in images. **Flow Cytometer experiments**: The cells were seeded in a six-well plate at a density of 5 × 10^5^ cells per well in respective medium. After 24 h, the cells were washed twice with 1 × PBS, and incubated with rhodamine B-labeled liposomes diluted with respective medium to a liposome concentration of 0.5 mg/mL for 4, 12 and 24 h at 37 ℃. The fluorescence intensity was measured with a BD LSRFortessa Cell Analyzer (BD, Franklin Lakes, USA). The mean fluorescence intensity of rhodamine B was measured with a BD LSRFortessa Cell Analyzer (BD, Franklin Lakes, USA). After excluding cellular debris and adherent cells by setting appropriate gates for Forward Scatter (FSC) and Side Scatter (SSC), the mean fluorescence intensity of rhodamine B was detected using the PE channel. A total of 10,000 events were collected for each sample.

### Cell uptake in co-culture model of HaCaT and MNT-1 cells

In order to evaluate the cell uptake of unmodified liposomes and HCMP-liposomes in the co-culture model of HaCaT and MNT-1 cells, Cell-Tracker Green CMFDA and Cell-Tracker Deep Red (Maokang Biotechnology Co., Ltd., Shanghai, China) were applied to HaCaT and MNT-1 cells respectively, prior to seeding. Cells were washed 3 times with 1 × PBS, then HaCaT cells was incubated with FBS-free medium containing Cell-Tracker Green CMFDA (5 *μ*M) and MNT-1 cells were incubated with FBS-free medium containing Cell-Tracker Deep Red (1 *μ*M) for 30 min at 37 °C. After incubation, the cells were washed 3 times with 1 × PBS. For CLSM observation, only HaCaT cells were labeled with Cell Tracker Green CMFDA, and then HaCaT and MNT-1 cells were cultured in a culture dish at a density of 10^5^ per well with the ratio of MNT-1 and HaCaT cells of 1:5 by using a CLSM as described above. For 2D flow cytometry experiments, HaCaT and MNT-1 cells were both labeled, and the cells were seeded in a six-well plate at a cell density of 5 × 10^5^ cells/well with the ratio of MNT-1 and HaCaT cells of 1:5. After 24 h, the cells were incubated with rhodamine B-labeled liposomes as described above. The fluorescence intensity was measured with a BD LSRFortessa Cell Analyzer (Ex/Em: Cell-Tracker Green CMFDA, 492/517 nm; Cell-Tracker Deep Red, 630/650 nm).

### Evaluation of melanosome transfer

By comparing the efficiency of melanosome transfer in co-culture models with different incubation conditions and seeding ratios of HaCaT and MNT-1 cells, Fetal Bovine Serum (FBS)-free medium and a 5:1 ratio were selected for evaluation of melanosome transfer (**Supplementary Fig. 8**). For CLSM observation of melanosome transfer in co-culture model, HaCaT and MNT-1 cells were culture at a culture dish at a density of 10^5^ per well and in a ratio of MNT-1 and HaCaT of 1:5 by using a Zeiss LSM880 confocal microscope. The culture media was composed of MNT-1 medium and HaCaT medium at 1:5 ratio. After 24 h, the cells were washed twice with 1 × PBS, and incubated with liposomes diluted with FBS-free medium to a liposome concentration of 0.5 mg/mL; for the group without liposomes, only FBS-free medium was added. After incubation, the co-culture models were harvested and washed with cold PBS, fixed in 4% paraformaldehyde for 10 min, washed with PBS containing 0.1% Triton-X100 for 5 min. MNT-1 cells and melanosomes were immune-stained with PMEL 17 Rabbit Monoclonal Antibody (Beyotime, Shanghai, China) and Alexa Fluor 488-labeled Goat Anti-Rabbit IgG (H+L) (Beyotime, Shanghai, China) and HaCaT cells were immune-stained with anti-pan cytokeratin mouse recombinant multiclonal antibody (Abcam, Cambridge, UK) and Alexa Fluor 647-labeled Goat Anti-Mouse IgG (H+L) (Beyotime, Shanghai, China). After immunofluorescence staining as described above, the cells were stained with DAPI Staining Solution for 4 min, and then imaged via CLSM (Ex/Em: Alexa Fluor 488-labeled Goat Anti-Rabbit IgG (H+L), 495/519 nm; Alexa Fluor 647-labeled Goat Anti-Mouse IgG (H+L), 651/667 nm; DAPI, 364/454 nm). ImageJ (National Institutes of Health) was used to quantify the number of PMEL-17-positive spots in per HaCaT cells in images.

To quantitatively assay melanosome transfer, stained cells were analyzed by a BD LSRFortessa Cell Analyzer. HaCaT and MNT-1 cells were co-cultured in a 6 well at a cell density of 5 × 10^5^ cells/well with the ratio of MNT-1 and HaCaT cells of 1:5. After 24h, the cells were incubated and immune-stained as described above. The fluorescence intensity was measured with a BD LSRFortessa Cell Analyzer. The number of HaCaT cells containing melanosomes were shown in the upper right quadrant (Q2), the melanosome transfer was quantified by analyzing the fluorescence intensity of Alexa Fluor 488 in the Q2 region.

### Isolation of pigment globules containing melanosomes from MNT-1 cell culture media

The collected MNT-1 culture media were centrifuged at 2000 × g for 10 min at room temperature (RT). The supernatants were centrifuged at 10000 × g for 30 min at RT. The precipitation was resuspended in MNT-1 cell culture media and centrifuged again in the same manner. After that, the precipitates were washed by PBS with brief and gentle mixing in order to avoid dispersion and were centrifuged again. Then the pellets were used as the pigment globule-rich fraction.

### Study on the mechanism of HCMP-liposomes interfering with melanosome transfer

Immunofluorescence was used to confirm that the cell secretions at the bottom of the cell culture dish and the cell secretions extracted from the culture medium were pigment globules containing melanosomes secreted by MNT-1 cells. MNT-1 cells were cultured in a culture dish at a density of 5 × 10^5^ per well, and the cells were immune-stained with PMEL 17 Rabbit Monoclonal Antibody and Alexa Fluor 488-labeled Goat Anti-Rabbit IgG (H+L); Isolated pigment globules were immune-stained with the same primary antibody and fluorochrome-labeled secondary antibody as described above. After Immunofluorescence, the cell secretions were stained with DAPI Staining Solution, and visualized via CLSM (Ex/Em: DAPI, 364/454 nm; Alexa Fluor 488-labeled Goat Anti-Rabbit IgG (H+L), 495/519 nm).

After proving that the cell secretions were pigment globules containing melanosomes, pigment globules at the bottom of cell culture dish and pigment globules isolated from cell culture medium were incubated with rhodamine B labeled liposomes to verify the interaction between liposomes and pigment globules. For pigment globules at the bottom of the cell culture dish, MNT-1 cells were cultured in a culture dish at a density of 5 × 10^5^ per well. After 24 h, the cells were washed twice with 1 × PBS, and incubated with rhodamine B labeled liposomes diluted with FBS-free medium to a liposome concentration of 0.5 mg/mL for 24 h at 37 ℃. After incubation, the pigment globules at the bottom of the cell culture dish were fixed with 4% Paraformaldehyde Fix Solution for 10 min, immune-stained with PMEL 17 Rabbit Monoclonal Antibody and Alexa Fluor 488-labeled Goat Anti-Rabbit IgG (H+L), stained with DAPI Staining Solution for 4 min, and then imaged via CLSM; For pigment globules isolated from cell culture medium, isolated pigment globules were incubated with rhodamine B labeled liposomes diluted with FBS-free medium to a liposome concentration of 0.5 mg/mL for 24 h at 37 ℃. After incubation, the isolated pigment globules were fixed with 4% Paraformaldehyde Fix Solution, immune-stained as described above, stained with DAPI Staining Solution, and then imaged via CLSM (Ex/Em: DAPI, 364/454 nm; Alexa Fluor 488-labeled Goat Anti-Rabbit IgG (H+L), 495/519 nm; Rhodamine B, 560/580 nm).

Cryo-TEM experiments were performed to confirm the adhesion of unmodified liposomes and HCMP-liposomes on the surface of pigment globules isolated from cell culture medium. In brief, the isolated pigment globules were suspended in 1 × PBS and incubated with liposomes at 37 ℃ for 24 h. After 24 h, the pigment globules were imaged by Cryo-TEM according to the method described above.

### Statistical analysis

The data were analyzed using GraphPad Prism (9.0.0). T-tests were performed to compare two sets of data, and one-way ANOVA was used to compare multiple sets of data. *P* < 0.05 was considered to indicate statistical significance. *P*-values correspond to \*\*\*\**P* ≤ 0.0001, \*\*\**P* ≤ 0.001, \*\**P* ≤ 0.01, \**P* ≤ 0.05, and ns: *P* ≥ 0.05.

## Acknowledgment

This research was financially supported by the Natural Science Foundation of China (22208121), Natural Science Foundation of Jiangsu Province (BK20221069), the Harvard MRSEC program of the NSF under Award No. DMR 20-11754 and the HealthInnoHK program of the Innovation and Technology Commission of the Hong Kong SAR Government. K.J. thanks the Alexander von Humboldt Foundation for financial support. Y.L. thanks the Postgraduate Research & Practice Innovation Program of Jiangsu Province (KYCX24_2541). We thank Harvard Center for Nanoscale Systems and Harvard Center for Biological Imaging for support.

## Contributions

K.J., D.A.W. and C.Y. supervised the work. C.L., Y.L. and K.J. designed the experiments. C.L. and K.J. operated the extrusion experiments. C.L. and Y.L. participated the microfluidics experiments. C.L. and Y.L. performed the cell experiments. Y.L. carried out the melanosome transfer experiments. K.J., C.L., D.A.W. and Y.L. wrote the manuscript. C.L., Y.L., C.X., C.Y., D.A.W. and K.J. participated in discussions. All authors contributed to the analysis and interpretation of the data.

