## Supplementary Information for "Engineering liposomes with cell membrane proteins to disrupt melanosome transfer"

#### Supplementary Methods

**Preparation of liposomes by extrusion method**

DOPC and cholesterol with chloroform were mixed at a molar ratio of 8:2 with 2.5 mM total lipid concentration. The mixture lipids were dried under a flow of nitrogen for 20 min and then vacuumed for 1h to from lipid film. The dried lipid film was hydrated with Phosphate Buffer Saline pH 7.4 (1x PBS) (Fisher Scientific), while vertexing. After hydration, the suspension was extruded eleven times through a polycarbonate filter of 0.1 µm using an Avanti Mini Extruder (Avanti Polar Lipids) at room temperature to obtain unmodified liposomes. Unmodified liposomes (2.5mM) were mixed with 0.2mg/ml HCMP at volume ratio of 10:1, 25:1, 50:1, 100:1, and 300:1, by using pipette up and down 5 times, to obtain the HCMP-liposomes with 6, 20, 30, 67 and 100 ug/mL HCMP concentrations, respectively.

**Cell uptake by A375 cells**: Liposomes were labeled with 18:1 Liss Rhod PE by adding 0.1 mol% 18:1 Liss Rhod PE in the lipids, followed by preparation procedures as described above. **Confocal observation experiments**: A375 cells were cultured in 24-well plate (ThermoFisher Scientific, Waltham, USA) density of 5 × $10^4$ per well for observation under a Zeiss LSM980 with incubation systems (CLSM, ZEISS, Oberkochen, Germany), the 20X objective were used. After 24 h, the cells were washed twice with 1 × PBS, and incubated with rhodamine-labeled liposomes for 24 h at 37 °C. After treatment, the cells imaged via CLSM (Ex/Em: rhodamine, 560/580 nm). ImageJ (National Institutes of Health) was used to quantify the relative fluorescence intensity of rhodamine in images. **Flow Cytometer experiments**: The cells were seeded in a 6 well at a density of 5 × $10^5$ cells per well in the medium. After 24 h, the cells were washed twice with 1 × PBS, and incubated with rhodamine-labeled liposomes diluted with respective medium for 24 h at 37 °C. The fluorescence intensity was measured with a BD LSRFortessa Cell Analyzer (BD, Franklin Lakes, USA). The mean fluorescence intensity of rhodamine was measured with a BD LSRFortessa Cell Analyzer (BD, Franklin Lakes, USA). After excluding cellular debris and adherent cells by setting appropriate gates for Forward Scatter (FSC) and Side Scatter (SSC), the mean fluorescence intensity of rhodamine was detected using the PE channel. A total of 10,000 events were collected for each sample.

### Supplementary Note

#### Theoretical calculations of the proportion of membrane protein covering the surface area of HCMP - liposomes

The projected area per phospholipid molecule in a bilayer membrane:

$S_{per\ phospholipid\ molecule} = 0.7\ nm^2$

Gram of phospholipid for HCMP: $m_{\mathrm{phospholipid\ of}\ HCMP} = 1.37\ mg$

The molecular weight of phospholipid: $\mathrm{M}_{\mathrm{phospholipid}} = 758.06\ g/mol$

The number of moles of phospholipid: $n_{phospholipid} = \frac{m_{\mathrm{phospholipid\ of}\ HCMP}}{\mathrm{M}_{\mathrm{phospholipid}}} = 1.81\ \mu mol$

The number of molecules of phospholipid: $N_{\mathrm{phospholipid}} = n_{phospholipid} \times N_A = 1.09 \times 10^{18}$

Membrane proteins incorporated into HCMP (1:300): $M_{HCMP\ (1:300)\ i} = 30.65\ \mu g$

Membrane proteins incorporated into HCMP (1:200): $M_{HCMP\ (1:200)\ i} = 43.17\ \mu g$

Membrane proteins incorporated into HCMP (1:100): $M_{HCMP\ (1:100)\ i} = 57.47\ \mu g$

The average molecular weight of HaCaT cell membrane proteins was assumed to be 100 kDa: $Mr_{HaCaT\ cell\ membrane\ proteins} = 100\ kDa$

The number of membrane protein moles of the HaCaT cell membrane protein on HCMP (1:300) surface:

$$n_{\mathrm{HaCaT\ cell\ membrane\ protein}\ HCMP\ (1:300)} = \frac{M_{HCMP\ (1:300)\ i}}{Mr_{HaCaT\ cell\ membrane\ proteins}} = \frac{30.65}{100,000} = 3.07 \times 10^{-4}\ \mu mol$$

The number of membrane protein molecules on HCMP (1:300) surface:

$$N_{\mathrm{HaCaT\ cell\ membrane\ protein}\ HCMP\ (1:300)} = N_A \times n_{\mathrm{HaCaT\ cell\ membrane\ protein}\ HCMP\ (1:300)} = 6.02 \times 10^{23} \times 3.07 \times 10^{-10} = 1.85 \times 10^{14}$$

The number of membrane protein moles of the HaCaT cell membrane protein on HCMP (1:200) surface:

$$n_{\mathrm{HaCaT\ cell\ membrane\ protein}\ HCMP\ (1:200)} = \frac{M_{HCMP\ (1:200)\ i}}{Mr_{HaCaT\ cell\ membrane\ proteins}} = \frac{43.17}{100,000} = 4.32 \times 10^{-4}\ \mu mol$$

The number of membrane protein molecules on HCMP (1:200) surface:

$$N_{\mathrm{HaCaT\ cell\ membrane\ protein}\ HCMP\ (1:200)} = N_A \times n_{\mathrm{HaCaT\ cell\ membrane\ protein}\ HCMP\ (1:200)} = 6.02 \times 10^{23} \times 4.32 \times 10^{-10} = 2.60 \times 10^{14}$$

The number of membrane protein moles of the HaCaT cell membrane protein on HCMP (1:100) surface:

$$n_{\mathrm{HaCaT\ cell\ membrane\ protein}\ HCMP\ (1:100)} = \frac{M_{HCMP\ (1:100)\ i}}{Mr_{HaCaT\ cell\ membrane\ proteins}} = \frac{57.47}{100,000} = 5.75 \times 10^{-4}\ \mu mol$$

The number of membrane peotein molecules on HCMP (1:100) surface:

$$N_{\mathrm{HaCaTcellmembraneprotein}HCMP(1:100)} = N_A \times n_{\mathrm{HaCaTcellmembraneprotein}HCMP(1:100)} = 6.02 \times 10^{23} \times 5.75 \times 10^{-10} = 3.46 \times 10^{14}$$

The average diameter of HaCaT cell membrane proteins was assumed to be 10 $nm$:

$D_{HaCaT\ cell\ membrane\ proteins} = 10\ nm$

The area of a single HaCaT cell membrane protein molecule is $\mathrm{S}_{\mathrm{single\ HaCaT\ cell\ membrane\ protein}}$:

$$\boldsymbol{S_{single\ HaCaT\ cell\ membrane\ protein\ molecule} = \pi(\frac{D_{HaCaT\ cell\ membrane\ proteins}}{2})^2 = 78.5\ nm^2}$$

The proportion of membrane protein covering the surface area of HCMP(1: 300)

$$= \frac{S_{single\ HaCaT\ cell\ membrane\ protein\ molecule} \times N_{\mathrm{HaCaT\ cell\ membrane\ protein}\ HCMP\ (1:300)}}{S_{per\ phospholipid\ molecule} \times \frac{N_{\mathrm{phospholipid}}}{2}} = 3.81\%$$

The proportion of membrane protein covering the surface area of HCMP(1: 200)

$$= \frac{\boldsymbol{S_{single\ HaCaT\ cell\ membrane\ protein\ molecule}} \times N_{\text{HaCaT cell membrane protein } HCMP \text{ (1:200)}}}{S_{per\ phospholipid\ molecule} \times \frac{N_{\text{phospholipid}}}{2}} = 5.35\%$$

The proportion of membrane protein covering the surface area of HCMP(1: 300)

$$= \frac{\boldsymbol{S_{single\ HaCaT\ cell\ membrane\ protein\ molecule}} \times N_{\text{HaCaT cell membrane protein } HCMP \text{ (1:300)}}}{S_{per\ phospholipid\ molecule} \times \frac{N_{\text{phospholipid}}}{2}} = 7.12\%$$

#### Supplementary Figures

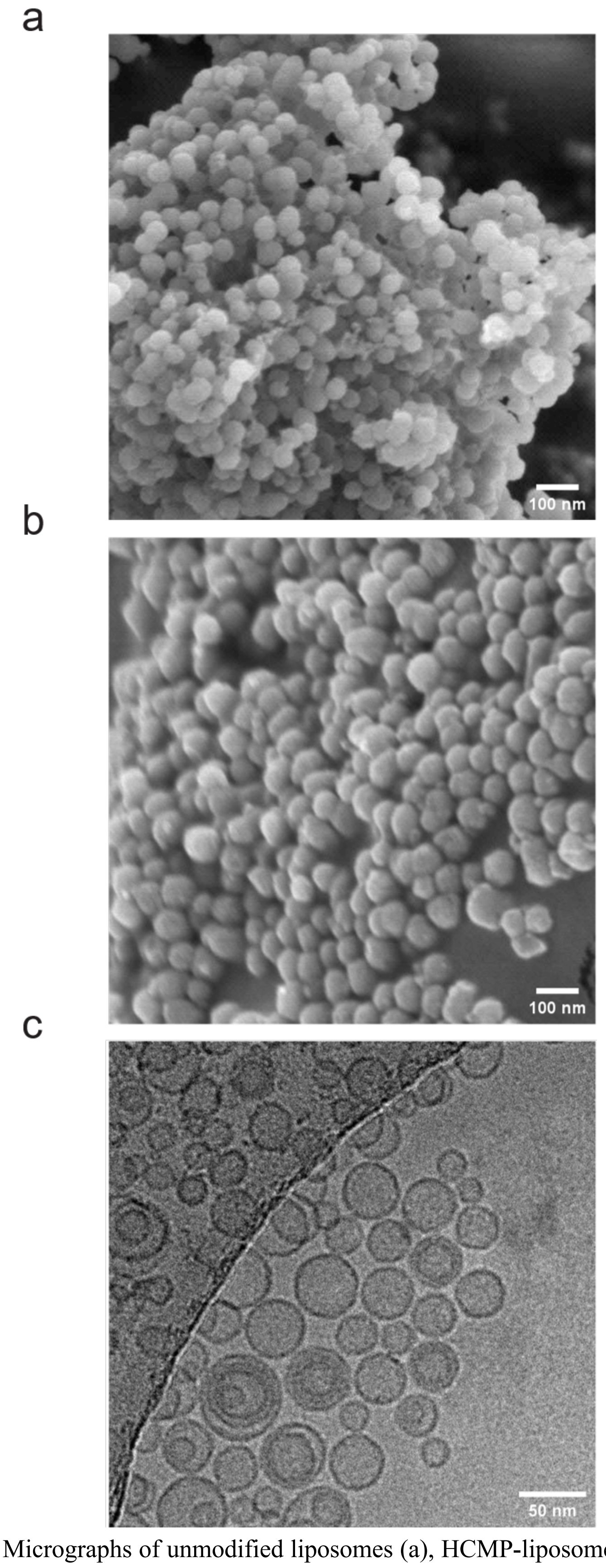


**Supplementary Fig. 1** Micrographs of unmodified liposomes (a), HCMP-liposomes (b) taken by SEM and micrographs of unmodified liposomes taken by Cryo-TEM (c).

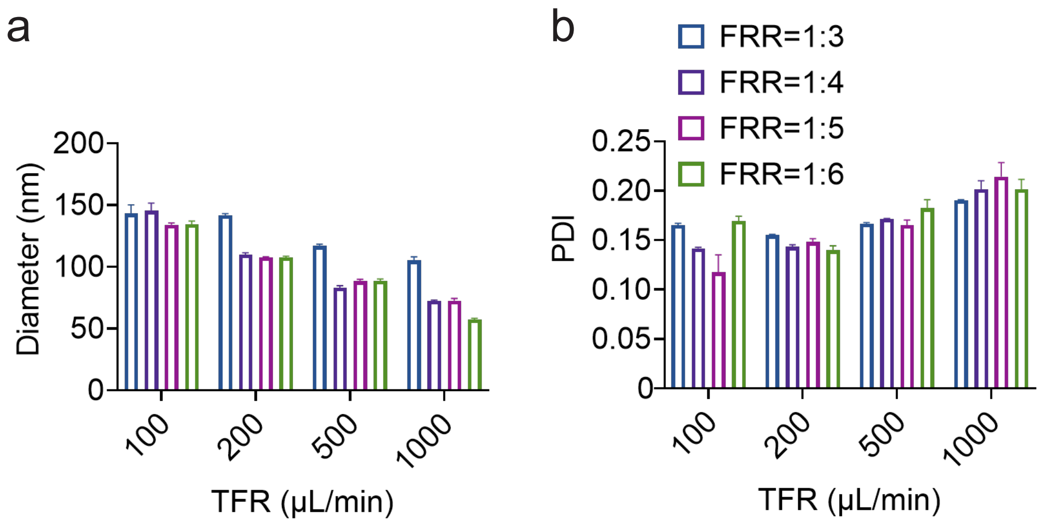


**Supplementary Fig. 2** The effects of the total flow rate (TFR) and flow rate ratio (FRR) of microfluidic chips on diameter (a) and PDI (b) of liposomes.

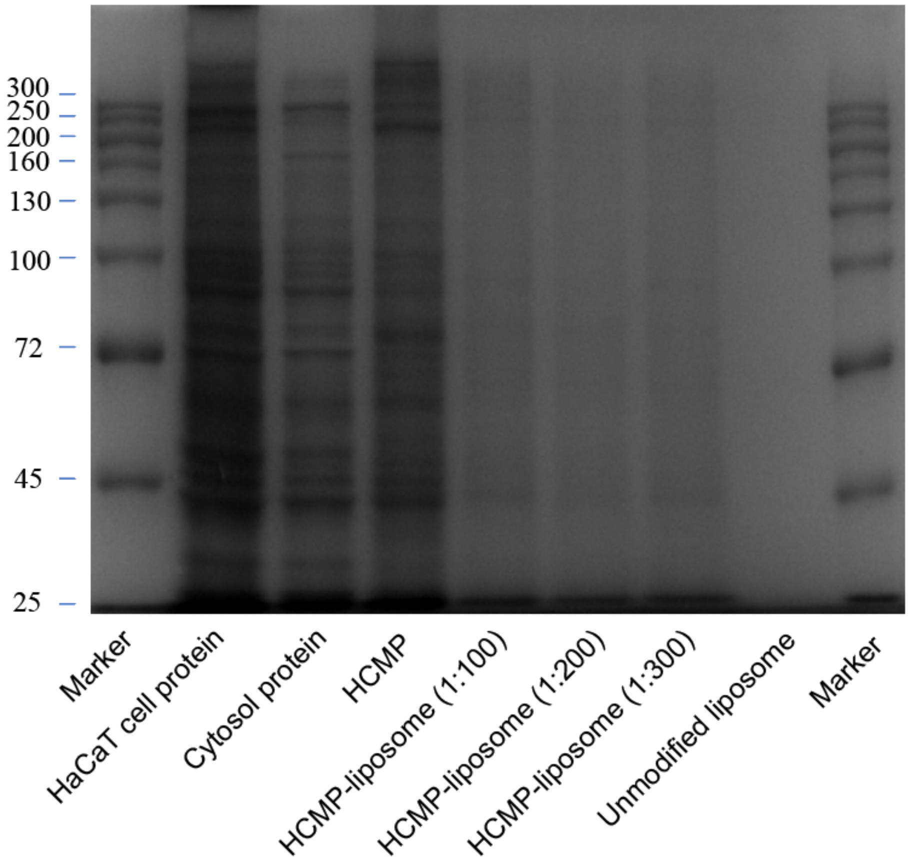


**Supplementary Fig. 3** HaCaT cell membrane proteins analyzed by SDS-PAGE.

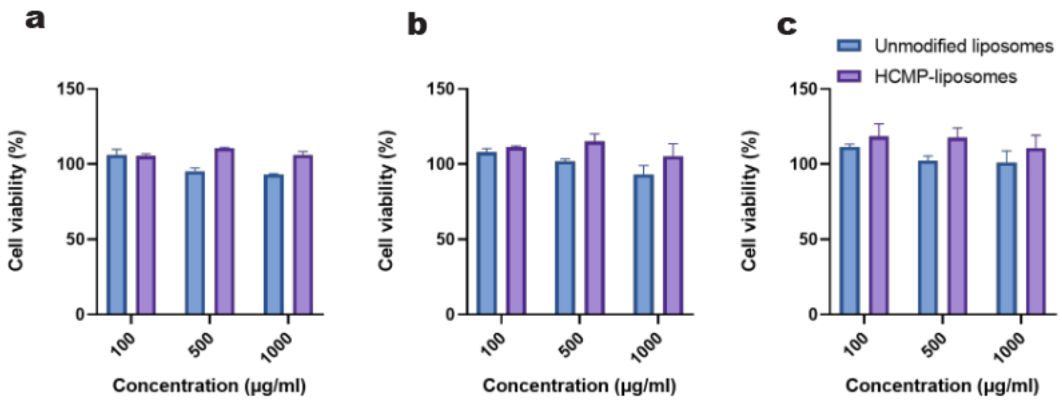


**Supplementary Fig. 4** Effects of unmodified liposomes and HCMP-liposomes on the viability of HaCaT cells (a), MNT-1 cells (b) and co-culture model (c), respectively.

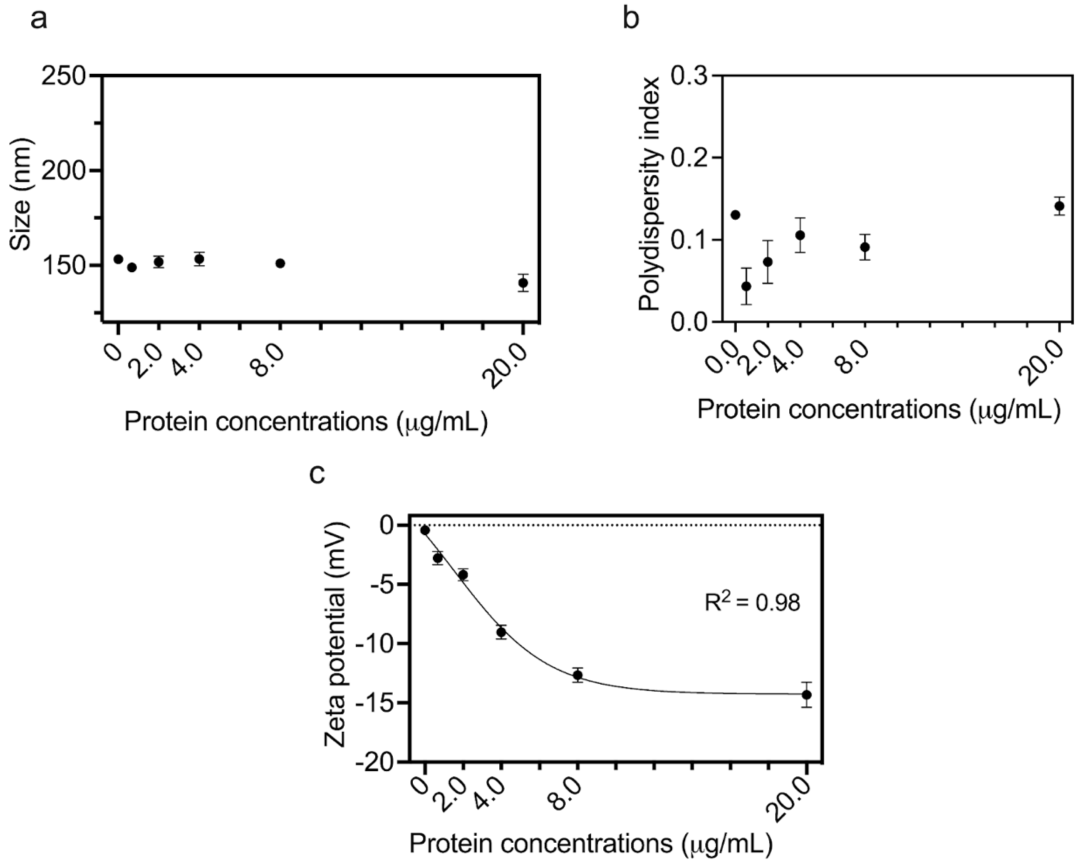


**Supplementary Fig. 5** The size (a), polydispersity index (b) and zeta potential (c) of HCMP-liposomes prepared with the extrusion method.

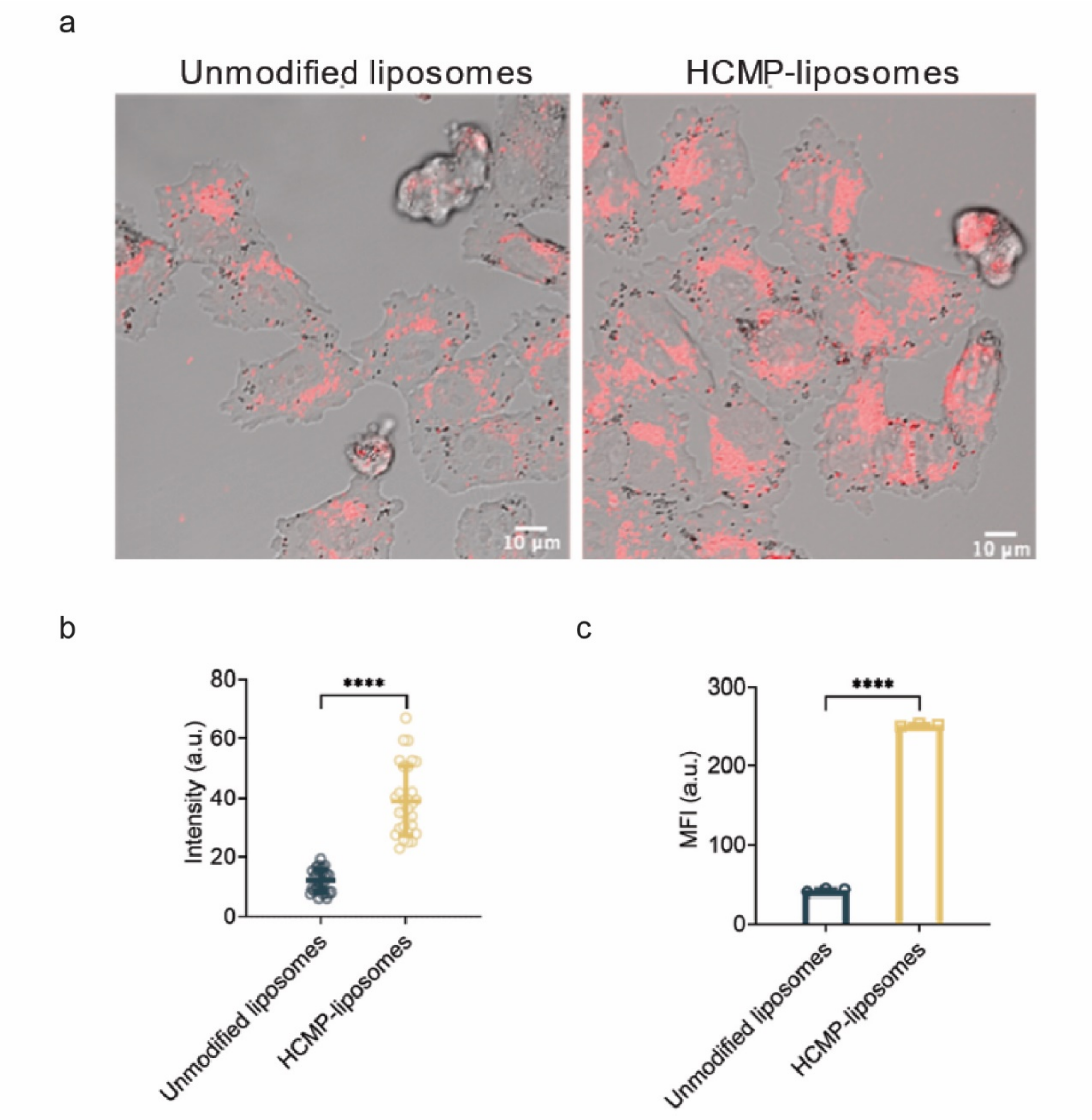


**Supplementary Fig. 6** Confocal images showing the A375 cell uptake of unmodified liposomes and HCMP-liposomes at 24 h (a) and fluorescence intensity analysis of cell uptake by A375 cells (n=30) (b). Flow cytometry profiles showing the A375 cell uptake of unmodified liposomes and HCMP-liposomes at 24 h (c).

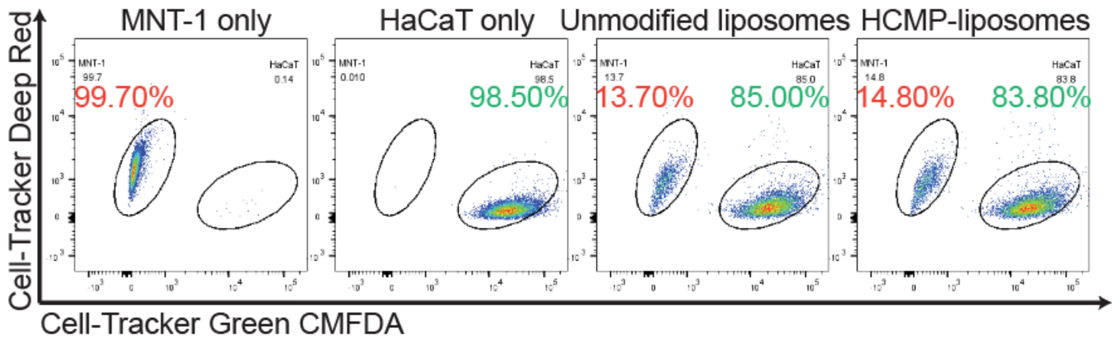


**Supplementary Fig. 7** The flow cytometry images of the HaCaT and MNT-1 cell uptake of unmodified liposomes and HCMP-liposomes in co-culture model after 24 h. HaCaT (Cell-Tracker Green CMFDA-labeled, $\lambda_{ex}$=492 nm) and MNT-1 cells (Cell-Tracker Deep Red-labeled, $\lambda_{ex}$=630 nm). The inserted values with red and green color represent the relative proportion of MNT-1 cells, respectively.

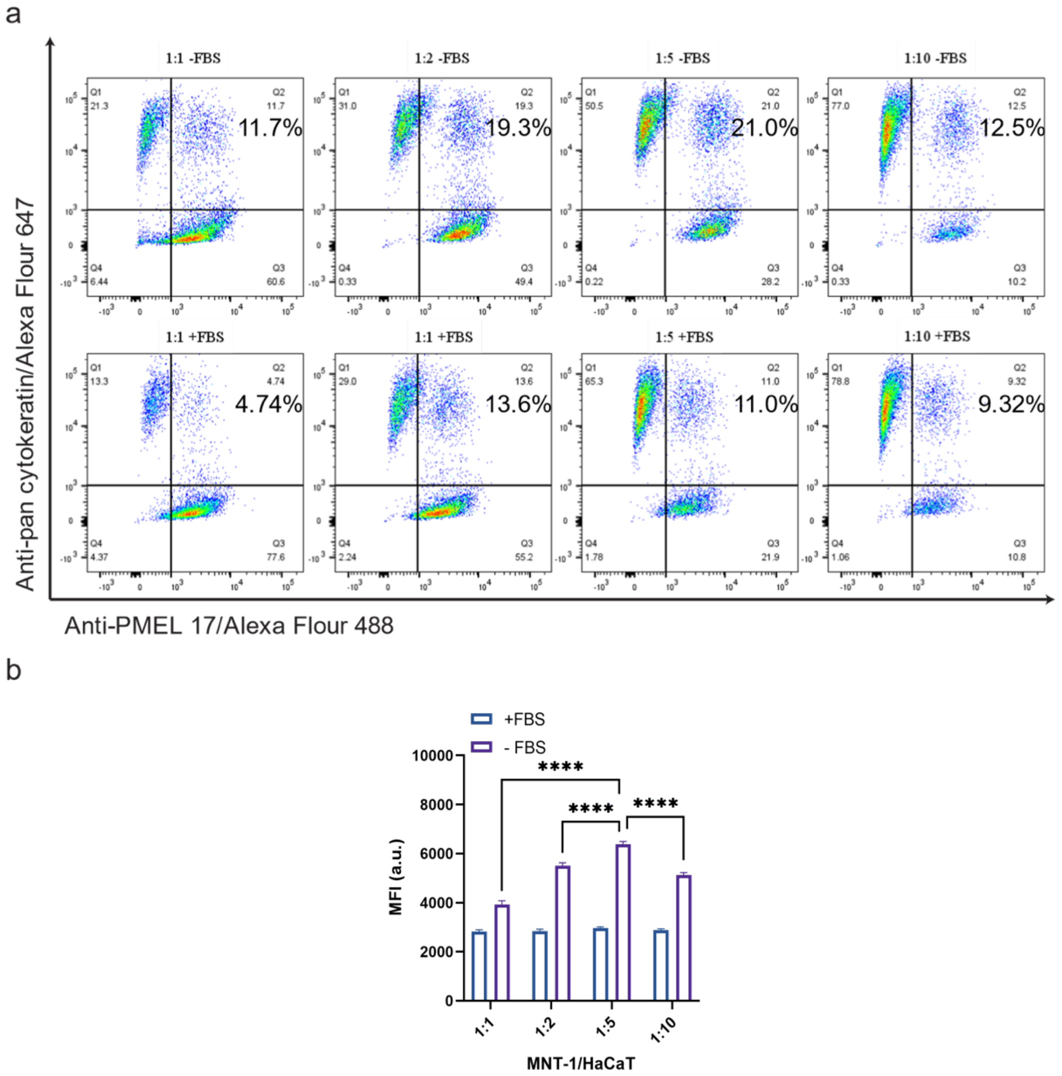


**Supplementary Fig. 8** Effects of cell ratio and media on melanosome transfer in co-culture model of HaCaT and MNT-1 cells. (a) Flow cytometry images of melanosomes transferred into HaCaT cell in co-culture model with ratio of MNT-1 and HaCaT cells of 1:1, 1:2, 1:5 and 1:10, and medium with and without FBS for 24 h. (b) Quantification analysis of mean fluorescence intensity of PMEL-17 in HaCaT cells (Q2 area) in the flow cytometry images.

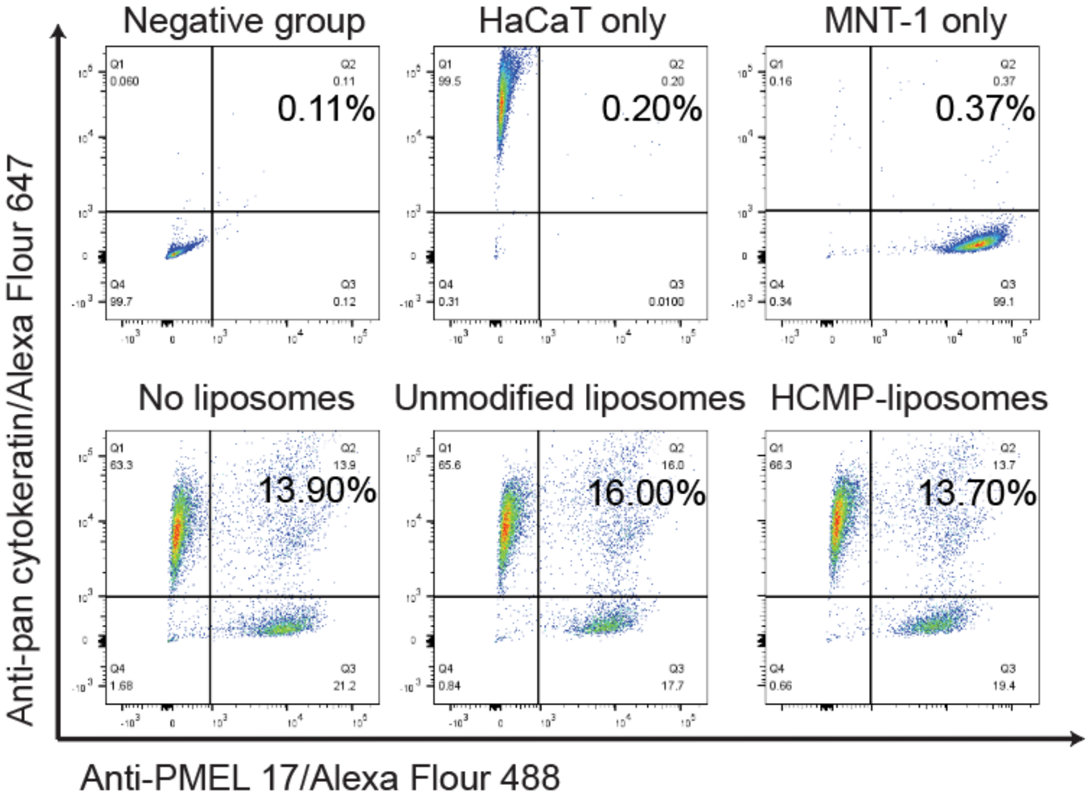


**Supplementary Fig. 9** Flow cytometry scatter plotsimages of melanosome transfer in co-culture model of HaCaT and MNT-1 cells. HaCaT cells immunostained with Anti-pan cytokeratin (($\lambda_{ex}$=651nm); MNT-1 cells and melanosomes immunostained with Anti-PMEL 17 (($\lambda_{ex}$=495nm). Q2 are expressed as the PMEL-17-positive spots in HaCaT cells. The inserted value represents the relative proportion of the Q2 area.

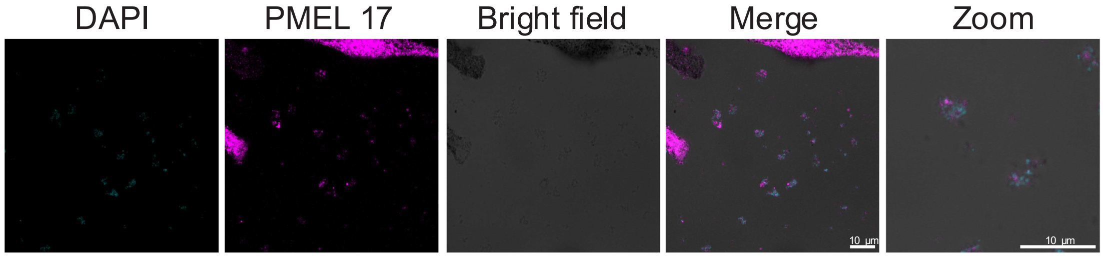


**Supplementary Fig. 10** CLSM of pigment globules in MNT-1 cells. Scale: 10 $\mu$m.

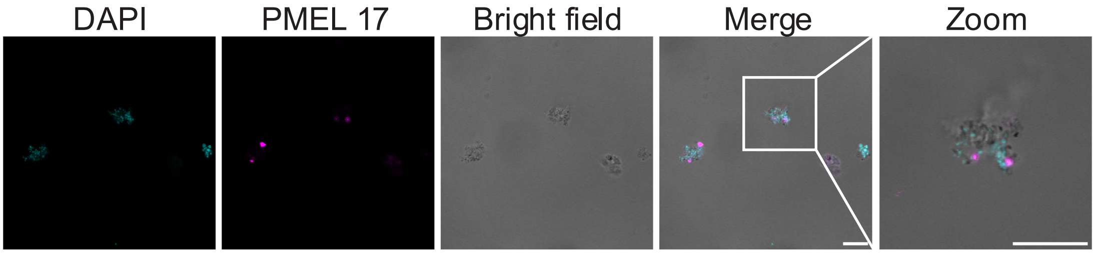


**Supplementary Fig. 11** CLSM of isolated pigment globules. Scale: 10 $\mu$m.

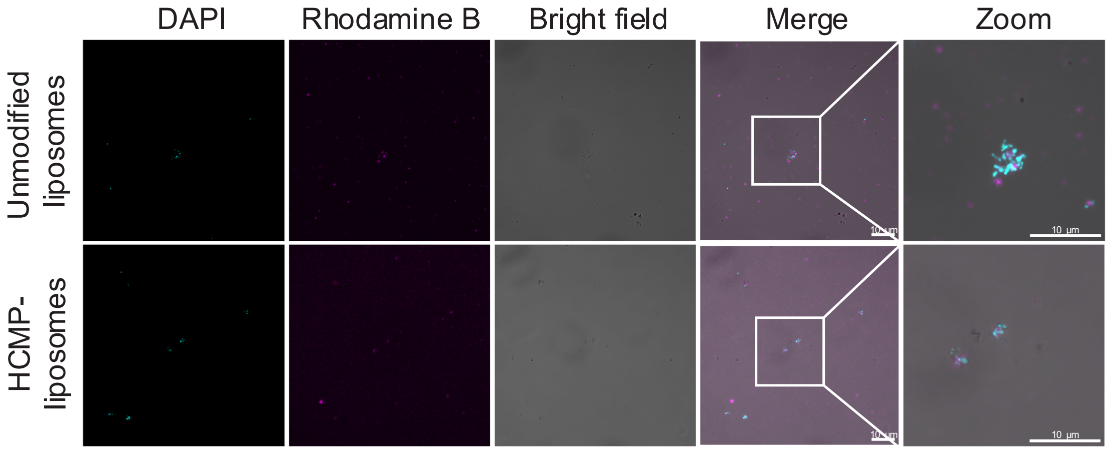


**Supplementary Fig. 12** CLSM of isolated pigment globules incubated with unmodified liposomes and HCMP-liposomes. Scale: 10 $\mu$m.

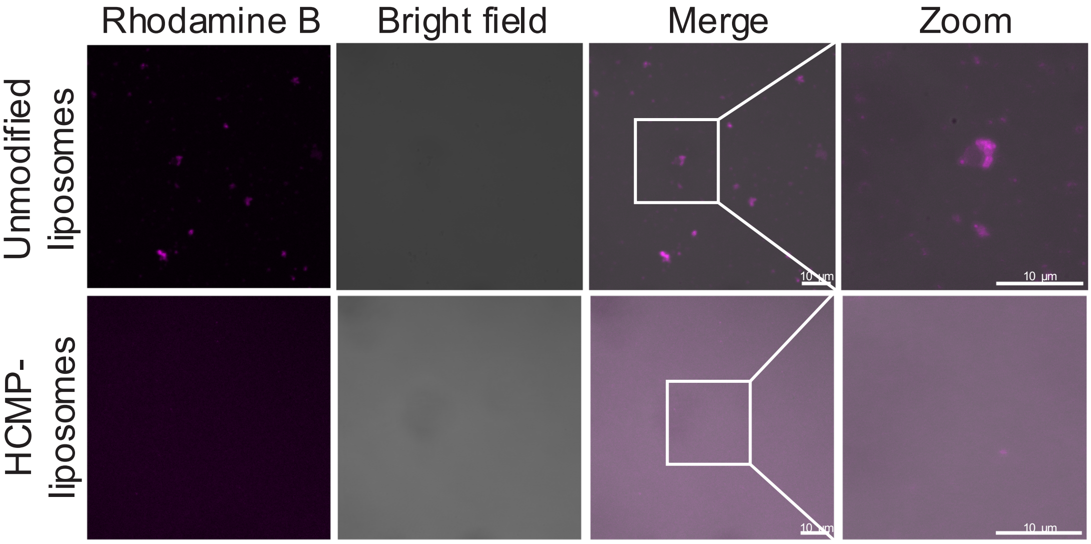


**Supplementary Fig. 13** CLSM of unmodified liposomes and HCMP-liposomes after incubation at 37 °C for 24 h. Scale: 10 $\mu$m.

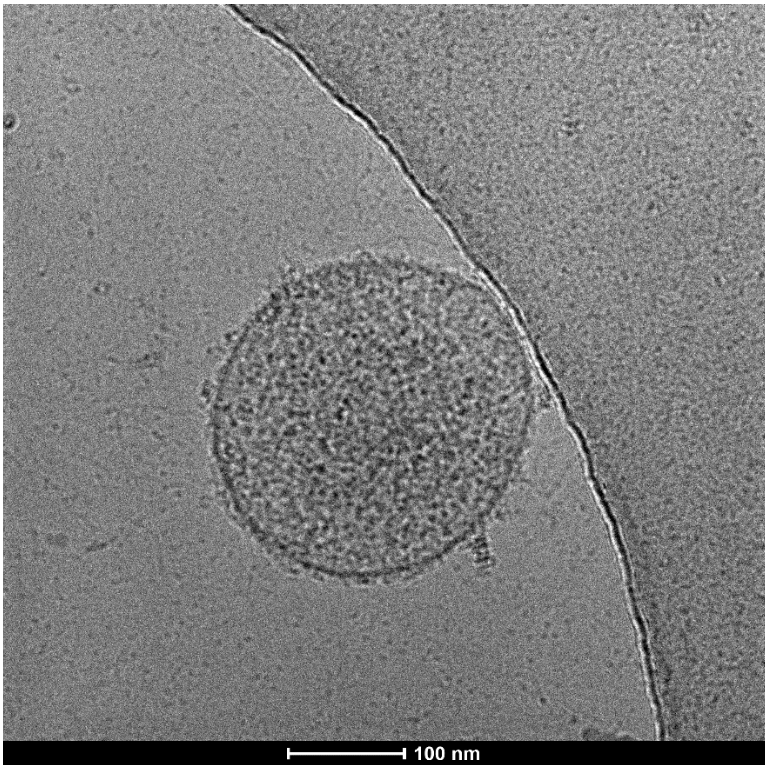


**Supplementary Fig. 14** Cryo-TEM of isolated pigment globules. Scale: 100 nm.
